# Humoral response to SARS-CoV-2 by healthy and sick dogs during COVID-19 pandemic in Spain

**DOI:** 10.1101/2020.09.22.308023

**Authors:** Ana Judith Perisé-Barrios, Beatriz Davinia Tomeo-Martín, Pablo Gómez-Ochoa, Pablo Delgado-Bonet, Pedro Plaza, Paula Palau-Concejo, Jorge González, Gustavo Ortiz-Diez, Antonio Meléndez-Lazo, Michaela Gentil, Javier García-Castro, Alicia Barbero-Fernández

**Affiliations:** Biomedical Research Unit, Universidad Alfonso X el Sabio, Avda Universidad 1, 28691 Villanueva de la Cañada (Madrid), Spain; Vetcorner, Calle Mosén José Bosqued 2, 50012 Zaragoza, Spain; ERVET-DIEZ BRU, Calle Tordesillas 4, 28925 Alcorcón (Madrid), Spain; Micros Veterinaria SL, Calle Profesor Pedro Cármenes, 24007 León, Spain; Faculty of Veterinary, Universidad Alfonso X el Sabio, Avda Universidad 1, 28691 Villanueva de la Cañada (Madrid), Spain; Laboklin GmbH & Co. KG, Steubenstraße 4, 97688 Bad Kissingen, Germany; Cellular Biotechnology Unit, Instituto de Salud Carlos III, Ctra Majadahonda-Pozuelo, Km 2, 28220 Majadahonda (Madrid), Spain

**Keywords:** SARS-CoV-2, dogs, mycoplasma, antibody, pneumonia

## Abstract

COVID-19 is a zoonotic disease originated by SARS-CoV-2. Infection of animals with SARS-CoV-2 are being reported during last months, and also an increase of severe lung pathologies in domestic dogs has been detected by veterinarians in Spain. Therefore it is necessary to describe the pathological processes in those animals that show symptoms similar to those described in humans affected by COVID-19. The potential for companion animals contributing to the continued human-to-human disease, infectivity, and community spread is an urgent issue to be considered.

Forty animals with pulmonary pathologies were studied by chest X-ray, ultrasound study, and computed tomography. Nasopharyngeal and rectal swab were analyzed to detect canine pathogens, including SARS-CoV-2. Twenty healthy dogs living in SARS-CoV-2 positive households were included. Immunoglobulin detection by different immunoassays was performed. Our findings show that sick dogs presented severe alveolar or interstitial pattern, with pulmonary opacity, parenchymal abnormalities, and bilateral lesions. Forty dogs were negative for SARS-CoV-2 but *Mycoplasma* spp. was detected in 26 of 33 dogs. Five healthy and one pathological dog presented IgG against SARS-CoV-2.

Here we report that despite detecting dogs with IgG α-SARS-CoV-2, we never obtained a positive RT-qPCR, not even in dogs with severe pulmonary disease; suggesting that even in the case of a canine infection transmission would be unlikely. Moreover, dogs living in COVID-19 positive households could have been more exposed to be infected during outbreaks.

## Introduction

We are currently in an international health emergency generated by the emerging zoonotic coronavirus SARS-CoV-2 that began its expansion by the end of the year 2019 in Wuhan (China) and has caused a pandemic in a few months.^1^ COVID-19 is a pathology with various clinical manifestations caused by SARS-CoV-2, and the severity of its infection is mainly associated with lung injury, with findings similar to macrophage activation syndrome that causes hyperinflammation and lung damage by an uncontrolled activation and proliferation of T lymphocytes and macrophages.^2,3^

Four genera of coronavirus have been described: Alphacoronavirus, Betacoronavirus, Gammacoronavirus, and Deltacoronavirus (α-CoV, β-CoV, γ-CoV, and δ-CoV) according to their genetic structure. With the detection of SARS-CoV-2 in humans, seven coronaviruses had been isolated from people, but only two of them (SARS-CoV and MERS-CoV) predominantly infect the lower airways and cause fatal pneumonia. Severe pneumonia is associated with rapid viral replication, massive inflammatory cell infiltration, and proinflammatory cytokine responses.^4^ In humans, the other coronaviruses cause mild upper respiratory tract infections in immunocompetent adults and serious symptoms in children and elderly people. α-CoV and β-CoV infect mammals and have been also described in dogs and cats; mostly they are responsible for respiratory infections in humans and gastroenteritis in animals. In dogs, canine enteric coronavirus (CCoV), an α-CoV, causes an enteritis of variable severity (rarely fatal) and develop immunity; however some of the recovered dogs become carriers with the ability to infect other dogs. However, the described fecal-oral transmission pattern for CCoV includes only the canine species, and is not currently postulated as a possible zoonotic agent. However, canine respiratory coronavirus (CRCoV), that belong to the β-CoV (like SARS-CoV-2), cause respiratory symptoms in dogs, in general with mild clinical signs^5^ and occasionally as a coinfection with other respiratory pathogens. Recently, the first cases of asymptomatic dogs infected with SARS-CoV-2 have been described.^6^

Due to the zoonotic origin of SARS-CoV-2 and the described transmission between species, the hypothesis of the spread between animals becomes more plausible.^7^ Cases of infected cats, dogs, tigers, lions, minks, and ferrets have been reported during SARS-CoV-2 outbreaks, and all of them had close contact with infected people.^6,8^ Experimental infections using different animal species report an increased susceptibility to infection by ferrets and cats, and indicate a low susceptibility to SARS-CoV-2 infection by golden hamsters, macaques, fruit bats, and dogs.^9^ Furthermore, under natural conditions, infections in more than 20 mink farms have been reported. The transmission pattern postulated is that virus was transmitted to the animals by infected people and then the virus spread between minks.^10^ Later, in May 2020, two workers of the mink farms that could have acquired the infection from the minks have been reported. These would be the first described transmissions of SARS CoV-2 from animal to human (apart the first one that originated the pandemic).^11^ Some data suggests that also transmission from minks to cats and dogs occurred in the farms.

The World Organization for Animal Health-OIE stated that some animals can become infected by being in permanent contact with infected people, although they note that there are no evidences defining the role of infected pets in the spread of SARS-CoV-2. To date, no cases of transmission from domestic or captive wild animals to humans have been described (excluding, if confirmed, mink farm workers). Information regarding the possibility of companion animals becoming infected is confusing and controversial. Some authors describe that dogs whose owners were positive for SARS-CoV-2, showed serological negative to SARS-CoV-2, postulating that pets are not virus carriers.^12^ By contrast, some cases of companion dogs have been reported that were positive by RT-qPCR detection^6^ and others that have developed neutralizing antibodies against SARS-CoV-2.^13^ Currently, around ten RT-qPCR-positive dogs have been detected worldwide (Hong Kong, Denmark, and USA, but none in Spain), all of them in close contact with COVID-19 positive humans.^14^ Less than half of them were asymptomatic, one presented mild respiratory illness, and only one present also neutralizing antibodies coursing with hemolytic anemia.^15^ On the other hand, two negative-PCR dogs have developed neutralizing antibodies, one was asymptomatic and the other had breathing problems, but it is not clear if those were related to the infection.^15^ Further, molecular testing of 3500 dogs, cat and horse companion animals were done by IDEXX Company in USA and Korea and no positive cases were found. No positive cases were reported in dogs exposed to SARS-CoV-2 in France.^16^ Another recent study in Italy carried out with pets has shown that none of the 817 animals studied was positive to SARS-CoV-2 by RT-qPCR test but 13 dogs and 6 cats had neutralizing antibodies.^13^

In this manuscript we report that, during the months of the pandemic, an increase of aggressive lung pathologies in dogs was detected by veterinarians in Spain. Therefore, it is important to determine the infectious agent and a potential role of a SARS-CoV-2 infection. Considering the information that is currently available, it is therefore necessary to better describe the pathological processes that could occur in those animals that could be infected by SARS-CoV-2 and also showing symptoms similar to those described in humans affected by COVID-19. It is also highly relevant to determine if dogs could become infected in a home environment where close human-pet relationships occur. Therefore, the potential for companion animals contributing to the continued human-to-human disease, infectivity, and community spread is an urgent issue to be considered. Here, we describe the study of sick and healthy dogs regarding a potential infection with SARS-CoV-2.

## Materials and Methods

### Clinical study

A prospective study with forty dogs presenting pneumonia was performed between April and June 2020 in Spain. A clinical follow-up of all patients was performed and mortality was also recorded. Twenty healthy dogs living with people affected by COVID-19 were included as animals exposed to the virus. Inclusion/exclusion criteria and more information are available in Suppl. Methods. The study was approved by the Ethical Committee of the Faculty of Health Sciences, Alfonso X el Sabio University and all dog owners gave written informed consent.

### Image analysis

Chest X-ray (CXR), thoracic radiographic and a study ultrasound were performed in sick dogs. The pattern type, distribution and intensity were analyzed. The pathological lung was recognized when the ultrasound lung rockets also called B lines were observed (Figure 1) or by the presence of other pulmonary ultrasound findings for consolidation (crushing, tissue, nodule sign) (Figure 1). Computed tomographic (CT) was perform to assess the lesions distribution, classified as generalized, focal, and uni or bilateral. More information is available in Suppl. Methods.

**Figure 1.**
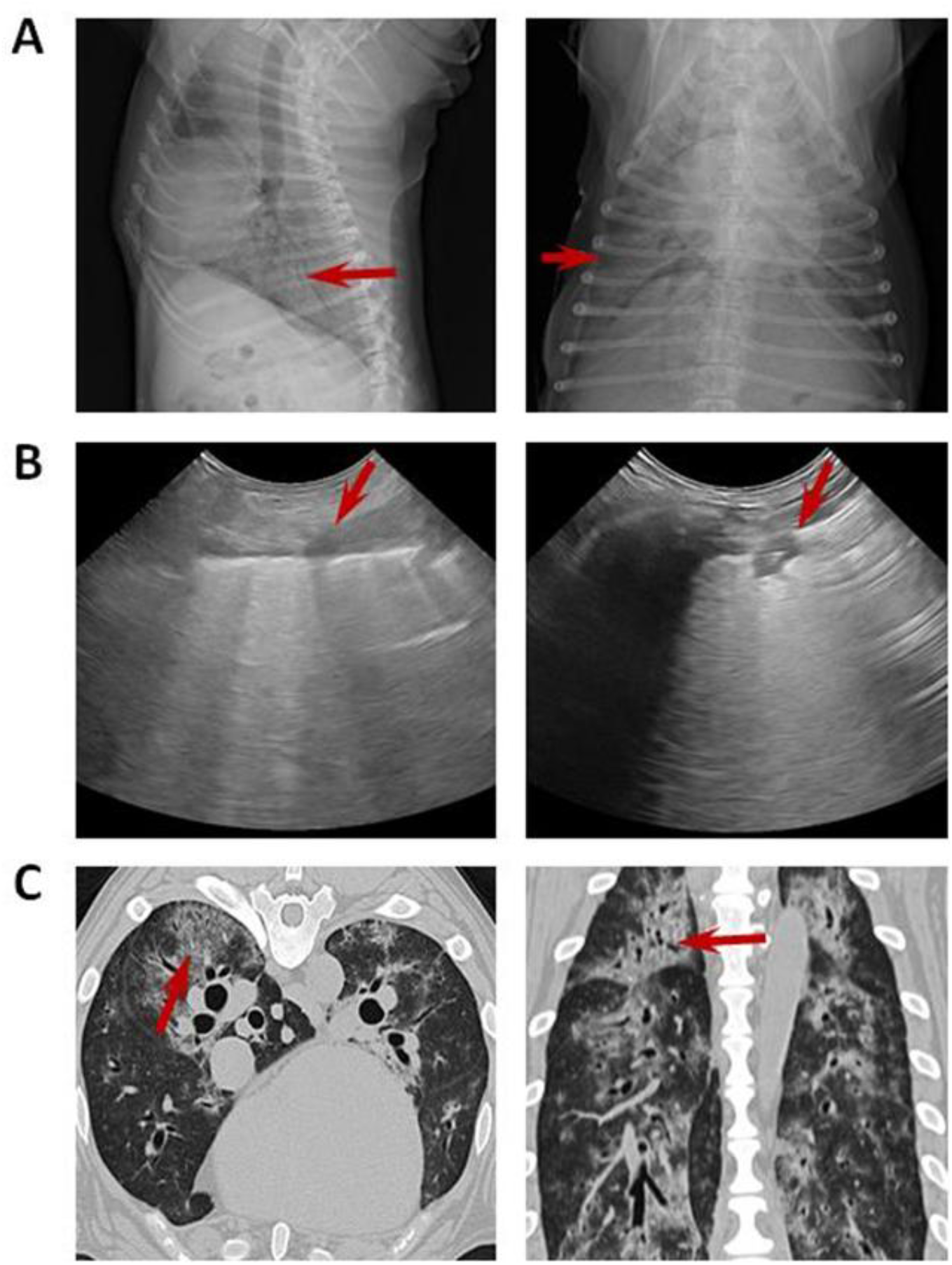
Imaging with chest radiograph, sonographic images and CT. (A) Thoracic radiograph made in right lateral (left) and dorsoventral (right) showing a generalized severe interstitial opacity accentuated in the caudodorsal (arrows). (B) Sonographic images of two patients with severe dyspnea showing a diffused B line (left; arrow) and consolidation focal lesions (right; arrow). (C) Transverse (left) chest CT images showing bilateral focal peripheral ground-glass opacities with intralobular and interlobular smooth septal thickening (arrow); sagital (right) chest CT images showing diffuse opacities with consolidation and bronchial wall thickening (arrow).

### Immunoglobulins detection by immunoassays

Blood samples from either sick or healthy-exposed dogs were analyzed to determine immunoglobulins (IgG) against SARS-CoV-2. A high-sensitive SARS-CoV-2 Spike S1 protein ELISA Kit was used. Values >2·5OD of the negative control were considered as positives. To determine antibodies (IgM and IgG) against CCoV, dog plasma samples were analyzed by EIA assays. To determine neutralizing antibodies (IgG) against canine adenovirus (CAV), canine parvovirus (CPV) and canine distemper virus (CDV), plasma samples were analyzed by an ELISA in solid phase. More information is available in Suppl. Methods.

### PCR analysis

Nasopharyngeal and rectal swabs were collected from sick dogs and analyzed by Laboklin GmbH & Co.KG using conventional PCR or real-time PCR (qPCR and RT-qPCR). All samples were tested for canine adenovirus type 2 (CAV-2), *Bordetella bronchiseptica*, CDV, canine parainfluenza virus (CPIV), canine influenza A virus (CIV), canine herpesvirus-1 (Canid alphaherpesvirus-1: CaHV-1), and SARS-CoV-2 by Taqman real-time PCR, and for *Mycoplasma* spp. by conventional PCR. More information is available in Suppl. Methods.

### Histopathology

Lungs of two dogs (SER209 and SER222) were histologically evaluated after necropsy. The macroscopic exam evaluated congestion, oedema and the lung injury pattern. Lung samples were fixed in formalin 4% for 24 hours, paraffin-embedded and 3µm thick sections stained with hematoxylin–eosin. More information is available in Suppl. Methods.

### Statistical methods

Categorical variables were presented as percentages. For continuous variables, data distribution normality was evaluated with the Kolmogorov-Smirnov test. Continuous data were presented as mean with standard deviation (SD) or median with interquartile range (IQR).

## Results

Forty pathologic dogs met the inclusion criteria with the mean age of 8 years (range: 2 months to 13 years). Fifteen breeds were recorded being the most common Cross-Breed 22·5% (9/40), Yorkshire terrier 12·5% (5/40), and German shepherd 10% (4/40). There were 22 females and 18 males.

The most common clinical signs were crackles on lung auscultation, followed by cough, tachypnoea, fatigue, fever, tachycardia, vomiting, and diarrhoea (Table 1). The radiographic findings in all analyzed dogs were consistent with mild to severe alveolar or interstitial pattern, with pulmonary opacity accentuated in the caudodorsal lung field. In 32·5% (13/40) of dogs a generalized increase in pulmonary opacity affecting all lung lobes was noted (Figure 1A). A focal alveolar infiltrate in the cranioventral lung field was detected in 50% (20/40) of dogs. One or more pathological findings were observed during the ultrasound examination of the patients. The main sonographic features of patients were dispersed B-line and rocket sign (100%; 31/31), partially diffused B-line (80·65%; 25/31), and pulmonary consolidations focal lesions (48·39%; 15/31) while complications as pleural effusion were rarely seen (4%; 1/25), further pneumothorax was not detected (Figure 1B). The CT findings were pulmonary parenchymal abnormalities (Figure 1C) and the lung lesions were bilateral distributed in all evaluated dogs.

**Table 1.** Epidemiologic characteristics, clinical features, and mortality

| Epidemiologic characteristics |  | Frequency<br>(N=40) | Percentage<br>(%) |
| --- | --- | --- | --- |
| Gender |  |  |  |
|  | Female | 22 | 55.0 |
|  | Male | 18 | 45.0 |
| Age (years) |  | 8.0* | 3.4** |
| Clinical features |  |  |  |
|  | Crackles (lung auscultation) | 40 | 100 |
|  | Cough | 37 | 92.5 |
|  | Tachypnoea (>30 breaths per minute) | 32 | 80.0 |
|  | Fatigue | 29 | 72.5 |
|  | Tachycardia (>130 beats per minute) | 24 | 60.0 |
|  | Fever (>39.5°C) | 23 | 57.5 |
|  | Vomiting | 15 | 37.5 |
|  | Diarrhoea | 14 | 35.0 |
| Mortality |  | 17 | 42.5 |

\* Mean \*\* Standard deviation

In order to assess the overall status of dogs, and considering that the number of white blood cells is frequently altered in an infection, we consider it relevant to perform a hematologic evaluation on twenty-four pathologic patients. The count of white blood cells was out of range in 58·3% of dogs being the number of neutrophils abnormal in 75%, lymphocytes in 37·5%, and monocytes in 45·8% (Table 2).

**Table 2.** Hematologic peripheral blood analysis. WBC: White Blood Cells.

| Hematologic | Out of range parameters | Normal | Mean | Standard deviation<br>(SD) |
| --- | --- | --- | --- | --- |
|  | N=24 (%) | range |  |  |
| WBC ($\times 10^3$ cells/μl) | 14 (58.3) | 6.0-17.0 | 20.5 | 9.7 |
| Neutrophils ($\times 10^3$ cells/μl) | 18 (75) | 3.0-11.5 | 16.7 | 9.1 |
| Lymphocytes ($\times 10^3$ cells/μl) | 9 (37.5) | 1.0-4.8 | 1.9* | 1.0-3.3** |
| Monocytes ($\times 10^3$ cells/μl) | 11 (45.8) | 0.2-1.4 | 1.2* | 0.7-1.7** |

\*Median \*\* Interquartile range (IQR)

Considering the altered number of immune cells observed in peripheral blood, together with the clinical course, we proceeded to evaluate possible pulmonary pathogens. In order to determine whether the observed pathologies could be related to a SARS-CoV-2 infection or to other pathogens, PCR analysis was performed. All forty dogs were negative for SARS-CoV-2 (Table 3). Furthermore, thirty-three dogs were analyzed for a complete profile including the most common canine infectious agents. All of them were negative for CPIV, CIV, and CaHV-1. *Mycoplasma* spp. and CDV were detected as a single agent with 57·6% (19/33) and 3% (1/33), respectively (Table 3).

**Table 3.** Infectious pathogens of sick dogs. CAV-2: canine adenovirus type 2; CDV: canine distemper virus; CPIV: canine parainfluenza virus; CIV: canine influenza A virus; CaHV-1: canine herpesvirus-1; SARS-CoV-2: severe acute respiratory syndrome coronavirus 2. Presence (+), absence (-), not determined (nd).

|  | CAV-2 | *Mycoplasma*<br>spp. | *Bordetella*<br>*bronchiseptica* | CDV | CPIV | CIV | CaHV-1 | SARS-CoV-2 |
| --- | --- | --- | --- | --- | --- | --- | --- | --- |
| SER 01 | - | + | + | - | - | - | - | - |
| SER 02 | - | + | - | - | - | - | - | - |
| SER 03 | - | + | - | - | - | - | - | - |
| SER 04 | nd | nd | nd | nd | nd | nd | nd | - |
| SER 05 | - | + | - | - | - | - | - | - |
| SER 06 | nd | nd | nd | nd | nd | nd | nd | - |
| SER 07 | nd | nd | nd | nd | nd | nd | nd | - |
| SER 08 | nd | nd | nd | nd | nd | nd | nd | - |
| SER 09 | - | + | - | - | - | - | - | - |
| SER 10 | - | + | - | - | - | - | - | - |
| SER 11 | - | + | - | - | - | - | - | - |
| SER 12 | - | + | - | - | - | - | - | - |
| SER 13 | - | + | - | - | - | - | - | - |
| SER 14 | - | + | - | - | - | - | - | - |
| SER 15 | - | + | - | + | - | - | - | - |
| SER 16 | - | + | - | - | - | - | - | - |
| SER 17 | - | + | - | - | - | - | - | - |
| SER 18 | - | + | - | + | - | - | - | - |
| SER 202 | - | - | - | - | - | - | - | - |
| SER 204 | - | + | - | - | - | - | - | - |
| SER 205 | - | + | - | - | - | - | - | - |
| SER 206 | - | + | - | - | - | - | - | - |
| SER 207 | - | + | - | - | - | - | - | - |
| SER 208 | - | + | - | - | - | - | - | - |
| SER 209 | - | + | - | - | - | - | - | - |
| SER 210 | - | + | - | - | - | - | - | - |
| SER 211 | - | - | - | - | - | - | - | - |
| SER 212 | - | - | - | - | - | - | - | - |
| SER 213 | - | - | - | - | - | - | - | - |
| SER 214 | + | + | - | - | - | - | - | - |
| SER 217 | - | + | + | + | - | - | - | - |
| SER 220 | - | + | - | + | - | - | - | - |
| SER 222 | - | + | - | - | - | - | - | - |
| SER 223 | - | - | - | - | - | - | - | - |
| SER 233 | - | - | - | + | - | - | - | - |
| SER 234 | - | + | - | + | - | - | - | - |
| SER 235 | - | - | - | - | - | - | - | - |
| SER 301 | nd | nd | nd | nd | nd | nd | nd | - |
| SER 302 | nd | nd | nd | nd | nd | nd | nd | - |
| SER 303 | nd | nd | nd | nd | nd | nd | nd | - |

The pathologies in our patients were very aggressive and 42·5% (17/40) of dogs died of pneumonia during follow-up (Table 1). Two of these deceased dogs have been necropsied in order to study lung tissue damage. The macroscopic exam show congested and oedematous, also consolidated lung parenchyma with patchy involvement in both dogs. Both animals presented severe interstitial pneumonia with diffuse alveolar damage. Furthermore, one dog (SER209) presented extensive congestion, haemorrhages (Figure 2A), and fibrin sheaths in the alveolar lumen with an inflammatory infiltrate of lymphocytes and macrophages (Figure 2B), and also scattered syncytia. Occasionally, alveoli are lined by Type II pneumocytes (Figure 2B). There was multifocal vasculitis with periarteriolar lymphocytes infiltrate and occasional vascular wall hyalinosis (Figura 2C). Animal SER222 showed intense autolytic changes with severe acute alveolar damage, vascular lesions such as congestion, rich-protein alveolar oedema (Figure 2D), and hyaline membranes that occluded alveolar lumina. Scant and disperse inflammatory cells, mainly macrophages with intracytoplasmic brown granular pigment were observed in alveolar septa. Interestingly, some of these findings, mainly the scattered syncytia, are usually present in some viral infections.^17^

**Figure 2.**
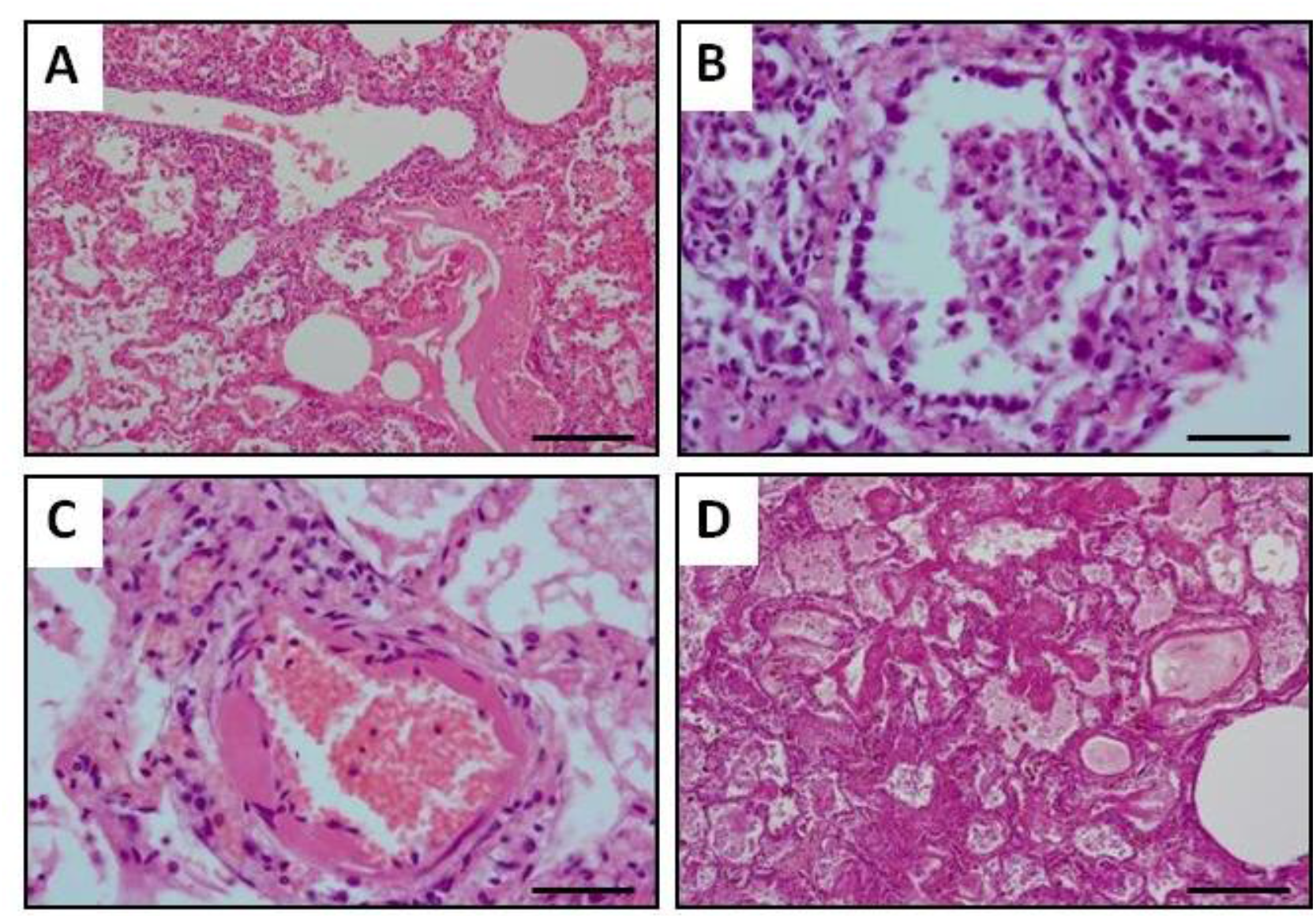
Histopathological study of lung tissues in pathologic dogs. Representative images of hematoxylin and eosin stained necropsy samples are shown. (A) Sample showing moderate vasculitis with rich-protein alveolar oedema and haemorrhages. (B) Lung tissue showing alveolar lined by type II pneumocytes and inflammatory infiltrate in the alveolar lumens. (C) Arteriolar wall hyalinosis is shown. (D) Diffuse alveolar damage with oedema and intralveolar hyaline membranes are shown. Scale bar: 200 µm (A and D) and 50 µm (B and C).

After analyzing the possible pathogens in the diagnosed dogs and the findings in the evaluated tissues, we set out to study the immune response against some of these infectious agents in 17 dogs. Further, we also decided to study 20 dogs that lived with people diagnosed with COVID-19, as a group of dogs exposed to the virus but which did not present symptoms at the time of sampling. First of all, information about vaccination was gathered to determine the immune status of dogs. Ten sick dogs had been vaccinated routinely as it is recommended by veterinarians but eight sick dogs did not received any vaccine (Suppl. Table S1). We have not detected any association between the vaccination patterns of pathological animals compared to healthy dogs.

Immunoglobulins G (IgG) against CAV, CPV, and CDV were analyzed in peripheral blood samples from these sick and healthy dogs (Table 4). Further, antibodies (IgM and IgG isotypes) against canine coronavirus that affects enteric tract, and also IgG against SARS-CoV-2 were studied in both groups (Table 4). The number of dogs that presented IgG antibodies against SARS-CoV-2 was higher in the group of healthy dogs (25%; 5/20), compared to the pathological ones (5·88%; 1/17). Interestingly, the sick dog that presented antibodies against SARS-CoV-2 was negative for the detection of the virus in swabs studied by RT-qPCR, however *Mycoplasma* spp. and CDV were detected in this patient (Table 3). All of the five IgG α-SARS-CoV-2 positives healthy dogs showed the same pattern of antibodies against the other studied pathogens, being positives for IgG α-CAV, IgG α-CPV, and IgG α-CDV (Table 4). Nevertheless, two of them presented IgG α-CCoV while the remaining three were not protected against canine coronavirus. Twelve healthy dogs presented IgG α-CCoV and two of them were positive for IgG α-SARS-CoV-2 (Table 4). Seven pathological dogs presented IgG α-CCoV but in this group all of them were negative to α-SARS-CoV-2 (Table 4).

**Table 4.** Immune response of sick and healthy dogs. IgG: immunoglobulin G; IgM: immunoglobulin M; CAV: canine adenovirus; CPV: canine parvovirus; CDV: canine distemper virus; CCoV: canine coronavirus; SARS-CoV-2: severe acute respiratory syndrome coronavirus 2. Presence (+), absence (-).

| | | IgG<br>α-CAV | IgG<br>α-CPV | IgG<br>α-CDV | IgM<br>α-CCoV | IgG<br>α-CCoV | IgG<br>α-SARS-CoV-2 |
| --- | --- | --- | --- | --- | --- | --- | --- |
| Pathologic dogs | SER 01 | + | + | + | + | + | - |
|  | SER 02 | + | + | + | - | + | - |
|  | SER 03 | nd | nd | nd | nd | nd | nd |
|  | SER 04 | + | - | + | - | - | - |
|  | SER 05 | + | + | + | - | + | - |
|  | SER 06 | + | + | + | - | - | - |
|  | SER 07 | + | + | + | - | - | - |
|  | SER 08 | + | + | - | - | - | - |
|  | SER 09 | + | + | + | + | + | - |
|  | SER 10 | + | + | + | - | + | - |
|  | SER 11 | - | + | - | - | - | - |
|  | SER 12 | + | + | + | - | - | - |
|  | SER 13 | + | + | + | - | - | - |
|  | SER 14 | + | + | + | + | - | - |
|  | SER 15 | - | + | + | - | - | + |
|  | SER 16 | + | + | + | - | - | - |
|  | SER 17 | + | + | + | - | + | - |
|  | SER 18 | + | + | + | - | + | - |
| Healthy dogs | SER 101 | + | + | + | - | - | + |
|  | SER 102 | + | + | + | - | - | + |
|  | SER 103 | + | + | + | - | - | - |
|  | SER 104 | + | + | - | - | + | - |
|  | SER 105 | + | + | + | - | + | - |
|  | SER 106 | - | - | + | - | + | - |
|  | SER 107 | + | + | + | - | - | - |
|  | SER 108 | + | + | - | - | + | - |
|  | SER 109 | - | + | - | - | - | - |
|  | SER 110 | + | + | + | - | + | + |
|  | SER 111 | + | + | + | - | + | - |
|  | SER 112 | + | + | + | - | + | - |
|  | SER 113 | + | + | + | - | + | - |
|  | SER 114 | + | + | + | - | + | - |
|  | SER 115 | + | + | + | - | - | - |
|  | SER 116 | + | + | + | - | + | - |
|  | SER 117 | + | + | + | - | + | + |
|  | SER 118 | + | + | + | - | + | - |
|  | SER 119 | + | + | + | + | - | - |
|  | SER 120 | + | + | + | - | - | + |

## Discussion

Professionals and pet owners demand more information about a possible infection of their animals with SARS-CoV-2. A survey of US veterinarians reported that 60% of them seeing owners concerned that their pets had COVID-19,^18^ so many owners are restless and demand a more in-depth clinical study from their animals. Pets are often in close contact with humans, and thus, it is important to determine their susceptibility to SARS-CoV-2 as well as the risk that infected pets are as a source of infection for humans.

Dogs are currently not considered to be susceptible hosts for SARS-CoV-2, despite few positive RT-qPCR test results in dogs were reported.^6^ Surveillance data from IDEXX laboratories, a multinational veterinary diagnostics company, showing that there were no positive results for SARS-CoV-2 in any of more than 1500 dog’s specimens submitted for respiratory PCR panels, according with the idea of transmission from human to pet is very rare. However, veterinarians, in Spain, have observed an increase in aggressive lung pathologies in dogs in the months of the human pandemic, which did not respond to conventional antibiotic treatments. Moreover, in veterinary medicine the respiratory disease is rarely lethal in the pet dog population. A mortality rate of 1·2% due to respiratory disease (only 0·3% due to pneumonia) has been reported,^19,20^ nevertheless in our study we found a mortality rate of 42·5% during follow-up without a clarified etiology. These dogs, with very aggressive lung diseases, showed a very similar appearance to those described for COVID-19 pneumonia in human medicine.^21^

Historically, the most common pathogens associated with canine infectious respiratory disease complex have been CPIV, CAV-2, *Bordetella bronchiseptica, Streptococcus equi* subsp. *zooepidemicus, Mycoplasma cynos*, CHV-1, CDV, CIV, and CRCoV.^20^ In our study, we detect eight of 33 analyzed dogs presenting classical primary respiratory pathogens, and we also detect IgM for CCoV in four dogs (3/17 pathological dogs and 1/20 healthy dog). In this regard, the presence of CRCoV is detected more frequently in dogs with mild clinical signs than in dogs with moderate or severe clinical signs,^5^ therefore, we would rule out that it was the agent responsible for the severe respiratory pathologies in these three dogs.

Interestingly 26 of 33 analyzed dogs showed a positive test for *Mycoplasma* spp. Many *Mycoplasma* spp. are commensal organisms that colonize the mucous membranes of the respiratory tract, and their role in canine infectious respiratory disease is not clear. Moreover, *Mycoplasma cynos* is the only *Mycoplasma* spp. significantly associated with pneumonia in dogs but it is still also unclear if *M. cynos* is a primary or secondary pathogen in dogs, because it can be cultured from the lungs of dogs, both with and without other identifiable infectious agents.^22^ In a European study of dogs with canine infectious respiratory disease seroprevalence of *Mycoplasma* spp. levels ranging from 20·7% to 61·9%,^23^ but in other study with healthy dogs mycoplasma were isolated from 78% to 93% of throat swabs.^20^ Moreover, mycoplasma infections are usually associated with other infections. It is interesting to note that mycoplasma coinfections are very common in COVID-19 human patients^24^ and it also has been suggested that a co-infection or activation of latent mycoplasma infections in COVID-19 disease may be important in determining a fatal disease course.^25,26^

Normally, the therapy response when treating respiratory tract diseases with drugs (antibiotics, bronchodilators, anti-inflammatories, antitussives, decongestants, mucolytics, mucokinetics or expectorants) is adequate or complete, nevertheless our patients did not respond adequately to the therapeutic protocol. A major pathogen has not been detected in our patients, so at the moment the causative agent of the pathologies is unknown. Further, the number of deaths was more than 30 times higher than expected without clarified etiology and curiously during the peak of the COVID-19 pandemic in Spain. When analyzing deceased dogs, interstitial pneumonia that usually courses with nonspecific lesions was detected, that has been described also in canine pathologies, such as canine distemper, herbicide poisoning or systemic processes (septicemia or uremia principally).^17^ However, it should be noted that showed lesions are similar to described for COVID-19 affected humans.^1^ Specially, striking lesions observed in vessels, both lymphocytic vasculitis, and the hyalinosis of the arteriolar wall.^27^ However, all of them were negative in RT-qPCR tests for SARS-CoV-2 using nasopharyngeal and rectal samples. These results agree with a large-scale study that recently has been shown to assess SARS-CoV-2 infection in 817 companion animals living in northern Italy and neither animals tested positive using RT-qPCR were found.^13^ Likewise, it should be considered that viral particles have been detected in the skin endothelium of human patients despite they were negative when tested by RT-qPCR. Therefore, it would be useful analyze SARS-CoV-2 by IHC in necropsy samples from our patients.^28^

Regarding the presence of immunoglobulins against SARS-CoV-2 in peripheral blood of pets, in a previous study 487 dogs were tested in China. They were serological negative for anti-SARS-CoV-2 IgGs. Among them, 15 pet dog and 99 street dog sera were collected from Wuhan City but it should be noted that only one pet dog living with a confirmed COVID-19 human patient presented antibodies against SARS-CoV-2.^12^ However, in the Italian study, 3·4% of 188 dogs (and 3·9% of 63 cats) had measurable SARS-CoV-2 neutralizing antibody titers. None of these animals with neutralizing antibodies displayed respiratory symptoms at the time of sampling. Interestingly, dogs from COVID-19 positive households seem to be significantly more likely to test IgG positive than those from COVID-19 negative households.^13^ Finally, it has been determined that only half of the dogs artificially inoculated with SARS-CoV-2 seroconverted.^8^

Here we detected specific anti-SARS-CoV-2 canine immunoglobulins in one sick dog (1/17) and in 25% (5/20) of dogs living in COVID-19 positive households, indicating their susceptibility to SARS-CoV-2 infection. In the Italian study they found 12·8% (6/47) of anti-SARS-CoV-2 canine immunoglobulins dogs in COVID-19 positive households, meanwhile only detected these immunoglobulins in 1·5% (2/133) living in COVID-19 negative households. Of note, in our study the influence of the family environment was not evaluated in the group of dogs with respiratory diseases. Some owners revealed that they had developed symptoms consistent with SARS-CoV-2 infection; nevertheless, they were not tested and confirmed. Other owners had not exhibited symptoms during the pandemic. Therefore, it was decided to exclude this data from the statistical study due to a lack of reliable information.

Most people infected with SARS-CoV-2 display an antibody response between day 10 and day 21 after infection, and several studies have suggested that previous antibodies and T cells against endemic human coronavirus may provide some degree of cross-protection to SARS-CoV-2 infection.^29^ Further, it has been reported pre-existing memory CD4+ T cells that are cross-reactive with SARS-CoV-2 and the common human cold coronaviruses HCoV-OC43, HCoV-229E, HCoV-NL63, or HCoV-HKU1.^30^ In our data we did not find any correlation between CCoV and SARS-CoV-2 IgG-positive dogs, although the low number of cases makes it difficult to reach a valid conclusion.

In sum, we analyzed dogs affected by severe pulmonary disease, all of them being negative for SARS-CoV-2 by RT-qPCR, however some of them present IgG α-SARS-CoV-2, as well as the healthy dogs; suggesting that even in the case of a canine infection it would be little transmissible. Moreover, dogs with owners positive for SARS-CoV-2 could have been more exposed to be infected during outbreaks.

## Acknowledgements

The authors would like to thank the owners of the dogs for their participation, as well as the veterinary clinics and hospitals that have collaborated in this study (Veterinary Hospital Vetcare, Veterinary Hospital Madrid Norte, among others). The authors would like to thank the “Centro de Transfusión Veterinario” for their donation of samples of virus not-exposed dogs.

