## Supplemental Information for "Humoral response to SARS-CoV-2 by healthy and sick dogs during COVID-19 pandemic in Spain"

### Supplemental Methods

#### Clinical study

This study was conducted in several hospitals and clinics, in Madrid and Zaragoza (Spain). Forty animals with pneumonia were enrolled to our study. The inclusion criteria were to present at least three of the following clinical signs: fever (rectal temperature higher than 39·5ºC), cough, fatigue, tachypnoea (higher than 30 breaths per minute), or crackles on lung auscultation. Gastrointestinal sings (vomiting and/or diarrhoea) and tachycardia (higher than 130 beats per minute) were also recorded. Dogs without evidences of pneumonia on imaging tests and dogs presenting signs that suggested cardiogenic oedema or tumors were excluded. The clinical diagnosis of pneumonia was through various imaging tests, thoracic ultrasound was performed. For a complete work-up dorsoventral and laterolateral thoracic radiographs, abdominal ultrasound, and complete hematologic work-up were performed on all dogs that owners authorized.

Healthy dogs did not present any symptoms at the time of taking samples, and therefore were considered healthy animals. The inclusion criterion was that the animals lived in homes where at least one person had been diagnosed with COVID-19. Exclusion criteria were pregnant females and dogs diagnosed with any ongoing pathology or infection.

#### Image analysis

Chest X-ray (CXR) was performed in two projections (Right Latero-Lateral (LL) /Dorsoventral (DV)). Thoracic radiographic images of dogs were independently reviewed by a veterinarian and were performed before the ultrasound study. The ultrasound study was performed using a portable ultrasonographic device (My Lab Alpha Vet Esaote S.P.A, Spain) equipped with two multifrequency linear (3-13 MHz) and microconvex (8-12 MHz) transducer. The Vet BLUE protocol was used to study the lungs of each patient, the images were acquired at 4 acoustic windows on each side of the thorax at standardized anatomic sites (caudal, perihilar, middle, and cranial), being a total of 8 sites/patient. Computed tomographic (CT) scans were obtained using 64-multidetector scanners (Aquilion, Toshiba) with all dogs positioned in sternal recumbency under general anaesthesia, the CT scan was performed during temporary apnoea induced by hyperventilation. CT scans were examined by a radiologist. The presence or absence of pleural effusion was studied with all techniques.

#### Peripheral blood analysis and immunoglobulins detection by immunoassays

Hematologic blood work was carried out in all dogs that owners authorized. Total white blood cells (WBC) were counted and individual populations of neutrophils, lymphocytes and monocytes were also recorded.

Blood samples to immunoassays were collected in BD vacutainer plasma separator tubes (BD PST II, Becton Dickinson), plasma was obtained and frozen at -20ºC until they were analyzed. As a negative control, samples of virus not-exposed dogs were used, kindly given by Centro de Transfusión Veterinario (Madrid, Spain). A high-sensitive SARS-CoV-2 Spike S1 protein ELISA Kit was used (MyBioSource) following the manufacturer's instructions. Captured IgG against SARS-CoV-2 S1 protein were detected by goat anti-dog IgG (H&L) polyclonal antibody conjugated with horse radish peroxidase (HRP) (Invitrogen). Absorbance was measured at 450nm using the Varioskan LUX, ver. 1.00.37 (Thermo Fisher Scientific). Results were calculated using the SkanIt Software 5.0 for Microplate Readers RE, ver. 5.0.0.42 (Thermo Fisher Scientific). Our cutoff value was 2·14OD to determine IgG against SARS-CoV-2. To determine IgM and IgG against CCoV, commercial EIA assays (Eurovet veterinaria S.L.) were used following the manufacturer's instructions. Absorbance was measured at 450nm using the Varioskan LUX, ver. 1.00.37 (Thermo Fisher Scientific). Results were calculated using the SkanIt Software 5.0 for Microplate Readers RE, ver. 5.0.0.42 (Thermo Fisher Scientific). The S/P ratio was calculated as (ODsample - MVODNC)/(MVODPC - MVODNC), where MV was mean value, NC was negative control and PC was positive control. When quantifying IgM, samples with the S/P ratio ≥0·250 were considered as positives. When quantifying IgG, samples with the S/P ratio ≥0·240 were considered as positives. ELISA in solid phase used was ImmunoComb Canine VacciCheck (Biogal Galed Laboratories Acs.). The assays were performed following the manufacturer´s instructions. ImmunoComb images were digitalized and spots densities were quantified using Image Lab™ 5.0 Software (Bio-Rad). Arbitrary units were calculated as follows: (sample spot intensity - sample mean background intensity) - (positive reference spot intensity - positive reference spot mean background intensity), and negative/positive criterion was applied following manufacturer's instructions.

**PCR analysis**

Swabs were incubated with lysis buffer for 1 hour at 65°C. Automated isolation of nucleic acids (RNA and DNA) was performed with the MagNA Pure 96 system (Roche diagnostics) according to manufacturer's instructions. Taqman real-time PCR was performed for CAV-2,[1] *Bordetella bronchiseptica,*[2] CDV,[3] CPIV, CIV,[4] canine herpesvirus-1,[5] and SARS-CoV-2[6] on a LightCycler®96 (Roche Diagnostics). Detection of *Mycoplasma* spp.[7] was performed by conventional PCR with subsequent gel electrophoresis. For each PCR run a negative and a positive control was included.

**Histopathology**

The lung injury pattern evaluates whether there is scattered involvement by areas or if it is generalized throughout the lungs, as well as the presence and type of lesions.

### Supplemental Tables

**Table S1.**

| | | CCoV | ICH | CPV | CDV | *Bordetella* | CPIV | *Leptospira* |
|---|---|---|---|---|---|---|---|---|
| Pathologic dogs | SER 01 | no | no | no | no | no | no | no |
| | SER 02 | yes | yes | yes | yes | yes | yes | yes |
| | SER 03 | yes | yes | yes | yes | yes | yes | yes |
| | SER 04 | yes | yes | yes | yes | yes | yes | yes |
| | SER 05 | yes | yes | yes | yes | yes | yes | yes |
| | SER 06 | no | no | no | no | no | no | no |
| | SER 07 | no | no | no | no | no | no | no |
| | SER 08 | yes | yes | yes | yes | yes | yes | yes |
| | SER 09 | yes | yes | yes | yes | yes | yes | yes |
| | SER 10 | yes | yes | yes | yes | yes | yes | yes |
| | SER 11 | no | no | no | no | no | no | no |
| | SER 12 | yes | yes | yes | yes | yes | yes | yes |
| | SER 13 | yes | yes | yes | yes | yes | yes | yes |
| | SER 14 | no | no | no | no | no | no | no |
| | SER 15 | no | no | no | no | no | no | no |
| | SER 16 | no | no | no | no | no | no | no |
| | SER 17 | yes | yes | yes | yes | yes | yes | yes |
| | SER 18 | no | no | no | no | no | no | no |
| Healthy dogs | SER 101 | no | yes | yes | yes | yes | no | yes |
| | SER 102 | no | yes | yes | yes | yes | no | yes |
| | SER 103 | no | yes | yes | yes | yes | yes | yes |
| | SER 104 | yes | yes | yes | yes | yes | yes | yes |
| | SER 105 | no | yes | yes | yes | yes | no | no |
| | SER 106 | yes | yes | yes | yes | dk/na | yes | yes |
| | SER 107 | no | yes | yes | yes | yes | yes | yes |
| | SER 108 | dk/na | yes | yes | yes | dk/na | dk/na | dk/na |
| | SER 109 | dk/na | yes | yes | yes | dk/na | dk/na | dk/na |
| | SER 110 | no | yes | yes | yes | yes | yes | yes |
| | SER 111 | no | yes | yes | yes | yes | yes | yes |
| | SER 112 | no | yes | yes | yes | yes | yes | yes |
| | SER 113 | no | yes | yes | yes | no | no | yes |
| | SER 114 | dk/na | yes | yes | yes | yes | yes | yes |
| | SER 115 | yes | yes | yes | yes | no | yes | yes |
| | SER 116 | yes | yes | yes | yes | no | yes | yes |
| | SER 117 | no | yes | yes | yes | yes | no | no |
| | SER 118 | yes | yes | yes | yes | yes | yes | yes |
| | SER 119 | dk/na | yes | yes | yes | yes | dk/na | yes |
| | SER 120 | dk/na | yes | yes | yes | yes | dk/na | yes |

**Table S1. Vaccination status from pathologic and healthy dogs.** Vaccines administered (yes, gray box); not administered (no, white box); don't know/no answer (dk/na). CCoV: canine coronavirus; ICH: infectious canine hepatitis; CPV: canine parvovirus; CDV: canine distemper virus. CPIV: canine parainfluenza virus.
